# Expanding the molecular and morphological diversity of Apusomonadida, a deep-branching group of gliding bacterivorous protists

**DOI:** 10.1101/2022.11.09.515670

**Authors:** Guifré Torruella, Luis Javier Galindo, David Moreira, Maria Ciobanu, Aaron A. Heiss, Naoji Yubuki, Eunsoo Kim, Purificación López-García

## Abstract

Apusomonads are cosmopolitan bacterivorous biflagellate protists usually gliding on freshwater and marine sediment or wet soils. These nanoflagellates form a sister lineage to opisthokonts and may have retained ancestral features helpful to understanding the early evolution of this large supergroup. Although molecular environmental analyses indicate that apusomonads are genetically diverse, few species have been described. Here, we morphologically characterize eleven new apusomonad strains. Based on molecular phylogenetic analyses of the rRNA gene operon, we describe four new strains of the known species *Multimonas media, Podomonas capensis, Apusomonas proboscidea* and *Apusomonas australiensis*, and rename *Thecamonas oxoniensis* as *Mylnikovia oxoniensis* n. gen., n. comb. Additionally, we describe four new genera and six new species: *Catacumbia lutetiensis* n. gen. n. sp., *Cavaliersmithia chaoae* n. gen. n. sp., *Singekia montserratensis* n. gen. n. sp., *Singekia franciliensis* n. gen. n. sp., *Karpovia croatica* n. gen. n. sp. and *Chelonemonas dolani* n. sp. Our comparative analysis suggests that apusomonad ancestor was a fusiform biflagellate with a dorsal pellicle, a plastic ventral surface and a sleeve covering the anterior flagellum, that thrived in marine, possibly oxygen-poor, environments. It likely had a complex cell cycle with dormant and multiple fission stages, and sex. Our results extend known apusomonad diversity, allow updating their taxonomy, and provide elements to understand early eukaryotic evolution.

---

APUSOMONADS are bacterivorous biflagellate eukaryotes found gliding on almost every kind of wet surface worldwide (see the detailed review by (Heiss et al., 2016). Although cosmopolitan, these nanoflagellate protists are not abundant (Larsen and Patterson, 1990; Patterson and Lee, 2000). Their discovery by classical optical microscopy approaches dates back to the beginning of the twentieth century. The first records of the fusiform *Amastigomonas* (Griessmann, 1913 [as *Rhynchomonas mutabilis*]; de Saedeleer, 1931) and the round *Apusomonas proboscidea* (Aléxéieff, 1924) reported gliding cells with a proboscis but no obvious flagella. Subsequently, by the end of the century, electron microscopy and ultrastructural observations of members of these two genera (e.g., Vickerman et al., 1974) led to their unification within the order Apusomonadida (Karpov and Mylnikov, 1989). This classification was notably based on the presence of a peculiar proboscis, consisting of a dynamic swinging sleeve enveloping the anterior flagellum, and an often hidden posterior flagellum trailing back underneath the ventral cell body (Karpov and Mylnikov, 1989). The characteristic morphology of these protists facilitated the observation and isolation of *Apusomonas proboscidea* on multiple occasions and that of the other described species of the genus, *Apusomonas australiensis* (Ekelund and Patterson, 1997). Likewise, several *Amastigomonas* species, with their characteristic fusiform shape, were described as well (Myl’nikov, 1999). The marine genus *Thecamonas*, morphologically similar, was segregated from this genus on the basis that, unlike *Amastigomonas*, its flagellum was clearly visible under light microscopy (Larsen and Patterson, 1990). Further ultrastructural studies of the cytology, the cytoskeleton and the flagellar apparatus of apusomonads (Karpov and Zhukov, 1986; Molina and Nerad, 1991; Karpov, 2007; Heiss et al., 2013) helped characterize the three-dimensional structure of these poorly studied protists, providing a deeper phenotypic basis to guide studies about early eukaryotic evolution.

The first molecular phylogenetic studies of apusomonads, using 18S rRNA genes as markers, confirmed the suspected monophyly of the group based on previous morphological observations (Cavalier-Smith and Chao, 1995; Cavalier-Smith and Chao, 2003). They also revealed their close evolutionary relationship with Opisthokonta, a large eukaryotic supergroup encompassing animals, fungi and their protistan relatives (Baldauf and Palmer, 1993; Cavalier-Smith and Chao, 1995; Kim et al., 2006), as apusomonads appeared as sister to opisthokonts, together with ancyromonads (Glucksman et al., 2011; Yubuki et al., in prep.). Accordingly, Cavalier-Smith forged the name Apusozoa to group both apusomonads and ancyromonads as sister to Opisthokonta (Cavalier-Smith, 2002). The monophyly of Apusozoa and its sister relation to Opisthokonta appeared supported by other early phylogenetic analyses based on 18S rRNA genes (e.g., Glucksman et al., 2011) and some housekeeping proteins (Paps et al., 2013). Subsequently, the position of apusomonads as sister to Opisthokonta has been consistently supported by larger phylogenomic analyses (Torruella et al., 2012; Derelle et al., 2015), often including the anaerobic breviates as closest relatives (Brown et al., 2013; Adl et al., 2019). However, many phylogenomic analyses repeatedly failed to retrieve the monophyly of ancyromonads and apusomonads, rendering Apusozoa paraphyletic (Brown et al., 2013; Cavalier-Smith et al., 2014; Torruella et al., 2015; Brown et al., 2018). Despite the uncertainty about their monophyly, Apusozoa appeared as a named group in databases such PR2 (Guillou et al., 2013), SILVA (Quast et al., 2013) and NCBI Taxonomy (https://www.ncbi.nlm.nih.gov/) for almost a decade. Accordingly, many environmental diversity studies put both groups of flagellates in the same bin to study biogeography and abundances (e.g., Sauvadet et al., 2010; Takishita et al., 2010), many of which may need revision in light of these more recent phylogenomic studies.

Regardless of their uncertain relationship with ancyromonads, apusomonads occupy a pivotal phylogenetic position from which to understand early eukaryotic evolution, as they seem to have retained some ancestral features. Among them is the biflagellate body plan (Cavalier-Smith and Chao, 2003) which, in its sister opisthokont lineage, evolved into the uniflagellate state inferred for animal and fungal ancestors (Torruella et al., 2018; Galindo et al., 2019; Galindo et al., 2021). Their phylogenetic interest led to sequencing the genome of *Thecamonas trahens* Larsen and Patterson, 1990 (Ruiz-Trillo et al., 2007). Interestingly, *Thecamonas* harbors genes that are related to multicellularity in animals (e.g., Sebe-Pedros et al., 2010). However, large phylogenomic studies are hampered by the limited taxon sampling available for the group. Currently, in addition to *Thecamonas*, the diversity of described apusomonads consists of a few genera and species including the fusiform *Amastigomonas*, which was subdivided into several genera (*Manchomonas*, *Podomonas, Multimonas* and *Chelonemonas*), and the round *Apusomonas* (Cavalier-Smith and Chao, 2010; Heiss et al., 2015). In 18S rRNA gene-based phylogenetic analyses, *Apusomonas* species branched within the clade of fusiform *Amastigomonas* (Cavalier-Smith and Chao, 2010; Heiss et al., 2015), making the latter genus paraphyletic. Accordingly, many of the organisms previously assigned to the genus *Amastigomonas*, including all currently cultured and sequenced examples, have been reassigned to new genera (Cavalier-Smith and Chao, 2010; Heiss et al., 2015). While no currently cultured apusomonad matches the original description of *Amastigomonas*, the historical use of the name suggests that the historically described *Amastigomonas* be taken in current context as a descriptive archetype, with no phylogenetic or taxonomic implications. Therefore, we use the term *’Amastigomonas-type‘* to refer to all apusomonads that lack the (probably derived) distinguishing characters of *Apusomonas*. As such, we apply this term to organisms representing notable morphological diversity and deep genetic divergence (Cavalier-Smith and Chao, 2010; Heiss et al., 2015; Heiss et al., 2016; Torruella et al., 2017). At any rate, despite the availability of only a handful of described species, environmental studies suggest a much wider apusomonad diversity (Torruella et al., 2017).

In this study, we aimed at expanding the diversity of described apusomonad species, updating their taxonomy, and gaining knowledge about their ecology and evolution. We cultured and investigated the morphology and molecular phylogeny of eleven new apusomonad strains isolated under aerobic conditions. Four of them are new strains of the known species *Multimonas media* Cavalier-Smith, 2010, *Podomonas capensis* Cavalier-Smith, 2010, *Apusomonas proboscidea* Aléxéieff, 1924 and *Apusomonas australiensis* Ekelund and Patterson, 1997. The remaining seven strains represent four new genera and six new species. We compare data from these new genera, species and strains with those of four other apusomonad species (*Podomonas magna* Cavalier-Smith, 2010, *Chelonemonas geobuk* Heiss et al., 2015, *Thecamonas trahens* Larsen and Patterson, 1990 and *Mylnikovia* [amended from *Thecamonas*] *oxoniensis* Cavalier-Smith, 2010).

## MATERIALS AND METHODS

### Sampling, isolation and culture conditions

Apusomonad strains used in this study were collected from sediment or soil from various marine and continental locations (Table 1). Natural samples were diluted with autoclaved marine or fresh water and let grow at room temperature until apusomonad cells were observed. Individual cells were either isolated through serial dilution or collected with an Eppendorf PatchMan NP2 micromanipulator using a 65 μm VacuTip microcapillary (Eppendorf) on an inverted microscope (Leica Dlll3000 B) and deposited in multiwell plates with marine or fresh water containing 1% YT medium (100 mg of yeast extract, 200mg of Tryptone in 100 ml of water: NIES, Japan) for growth. Strains were subcultured every two weeks and maintained at 15°C. Culture aliquots containing 10% DMSO were rapidly frozen and subsequently maintained in liquid nitrogen. They are available upon request from the culture collection of the DEEM laboratory, CNRS and Université Paris-Saclay (France). To acclimate *Cavaliersmithia chaoae* strain FABANU to freshwater, cells were subcultured every three days in marine water progressively diluted to freshwater (steps of 10% less marine water content). All strains were cultured under aerobic conditions.

**Table 1.** Isolates and sampling sites for cultured apusomonad strains investigated in this study. Previously described isolates are marked with an asterisk (*). Metadata of CCAP 1901/2, 1901/4 is taken from Cavalier-Smith and Chao (2010), KM053 are taken from Heiss et al. (2015), and ATCC 50062 from Larsen and Patterson (1990).

| Taxonomic name | Isolate | Sampling site | Sampling environment | Accession number |
| --- | --- | --- | --- | --- |
| *Multimonas media* | MMROSKO2018 | Roc'h ar Bleiz, Roscoff, France | Intertidal sediment | OM966643 |
| *Podomonas magna* | JJP-2003* | Kartesh, Baltic Sea, White Sea, Russia | littoral sample | OM966636 |
| *Podomonas capensis* | SPRINTER | New Jersey, USA | beach sediment | OM966641 |
| *Mylnikovia* (formerly *Thecamonas*) *oxoniensis* | IVY8c* | Radcliffe Square, Oxford, England | surface of ivy leaf | OM966635 |
| *Catacumbia lutetiensis* | ORSAYFEB19A PU2 | Orsay University campus, France | street gutter | OM966639 |
| *Catacumbia lutetiensis* | CAT | Paris Catacombs, France | subterranean water | OM966640 |
| *Cavaliersmithia chaoae* | FABANU | Menton, Provença, France | *Neorhodella* CCAP 1346/1 | OM966645 |
| *Apusomonas australiensis* | PROMEX | Tropical forest, Mexico | dry sediment | OM966637 |
| *Apusomonas proboscidea* | MPSANABRIA15 | Sanabria lake, Zamora, Spain | sediment | OM966642 |
| *Singekia franciliensis* | ORSAYFEB19A PU1 | Orsay University campus, France | street gutter | OM966638 |
| *Singekia montserratensis* | MONTSE19P8 | Montserrat Mountain, Catalonia, Spain | Puddle of rainwater | OM966647 |
| *Karpovia croatica* | CRO19P6 | Malo jezero, Mljet island, Croatia | sediment of saltwater lake | OM966646 |
| *Chelonemonas dolani* | D513M | Villefranche Bay, France | Mesopelagic water column (-250m) | OM966634 |
| *Chelonemonas geobuk* | KM053* | Masan Bay, Changwon, Korea | Intertidal sediment | OM966634 |
| *Thecamonas trahens* | 202 35m SS* | Sargasso Sea, Atlantic Ocean | marine | NW_013657636 |

### PCR amplification and sequencing of 18S rRNA genes and rRNA operons

For the molecular identification of apusomonads during preliminary steps, 40 μl of enrichment cultures were centrifuged in an Eppendorf tube at 13,400 rpm for 15 min at room temperature. The supernatant was removed until ~4 μl remained; cells were homogenized in that volume and lysed by applying two 5-second microwave pulses. This suspension was used as template DNA material to amplify the 18S rRNA gene applying a nested PCR protocol. First, the general eukaryotic primers 82F (5’-GAAACTGCGAATGGCTC) and 1520R (5’-CYGCAGGTTCACCTAC) were used. Then, a second PCR was carried out using 1 μl of the first PCR reaction as a template with the general eukaryotic primers 612F (5’-GCAGTTAAAAAGCTCGTAGT) and 1498R (5’-CACCTACGGAAACCTTGTTA). Both PCR reactions consisted of an initial denaturing period (95°C for 3 min), followed by 35 cycles consisting of three steps (denaturing, 93°C for 45 s; annealing, 5 cycles at 45°C and 30 cycles at 55°C, for 45 s; extension, 72°C for 2 min), and a final elongation period (72°C for 5 min). Amplicons were sequenced using the Sanger method (Genewiz, Leibniz, Germany). These 18S rRNA gene short amplicons were used for initial identification of apusomonads and for the selection of strains to be further analyzed. Subsequently, the full rRNA operons of selected strains were retrieved from RNAseq assemblies (Table 1). Briefly, total RNA was purified from apusomonad cultures using an RNAeasy kit (Qiagen, Germany). Eukaryotic RNA was sequenced after poly(A) selection by Eurofins (Konstanz, Germany). After preliminary assembly (Torruella et al., in prep.), contigs containing rRNA operons were identified by BLAST (BLASTn v2.2.26+; Altschul et al., 1990) using the corresponding 18S rRNA gene amplicon sequences as query.

### Sequence and phylogenetic analyses

For proper phylogenetic analyses, we used a total of 14 new apusomonad full rRNA operons from 11 new strains retrieved from preliminary RNAseq assemblies (Table 1). These were added to a previous 18S rRNA gene alignment covering the diversity of cultured and environmental apusomonads (Torruella et al. 2017) and aligned using MAFFT (Katoh et al., 2019) with the ‘add whole sequence’ feature. The alignment contained sequences of 139 apusomonads, 11 breviates and three environmental sequences of the NAMAKO environmental clade, which placed as sister to these groups in previous phylogenetic analyses, and were recognized as similar in the SILVA database (Quast et al., 2013). Spurious and unambiguously aligned sites were manually trimmed using AliView (Larsson, 2014). The resulting multiple sequence alignment (MSA) contained 1,877 nucleotides. It was used to infer a phylogenetic tree by maximum likelihood (ML) with IQ-TREE v1.6.11 (Nguyen et al., 2015) with the GTR+F+R4 evolutionary model. Node support was evaluated by 1,000 non-parametric bootstrap replicates (Felsenstein, 1985) (Figure 1). In addition to the 14 full rRNA operons sequenced in our study (Table 1), we retrieved that from *Pygsuia biforma* (Brown et al., 2013) after blasting (BLASTn v2.2.26+; Altschul et al., 1990) the corresponding region from *Thecamonas trahens* ATCC 50062 (NW_013657636) against the GenBank nr database. These gene operons were aligned using the MAFFT E-INS-i iterative method and trimmed with trimAl (Capella-Gutierrez et al., 2009). This MSA, containing 16 sequences and 5,596 nucleotide positions, was used to infer an ML phylogenetic tree with the IQ-TREE webserver (Trifinopoulos et al., 2016) using the GTR+F+R4 model and 1,000 nonparametric bootstraps (Figure 2). All data files are available at figshare (10.6084/m9.figshare.19319594). Sequences have been deposited in GenBank (accession numbers given in Table 1).

**Figure 1.**
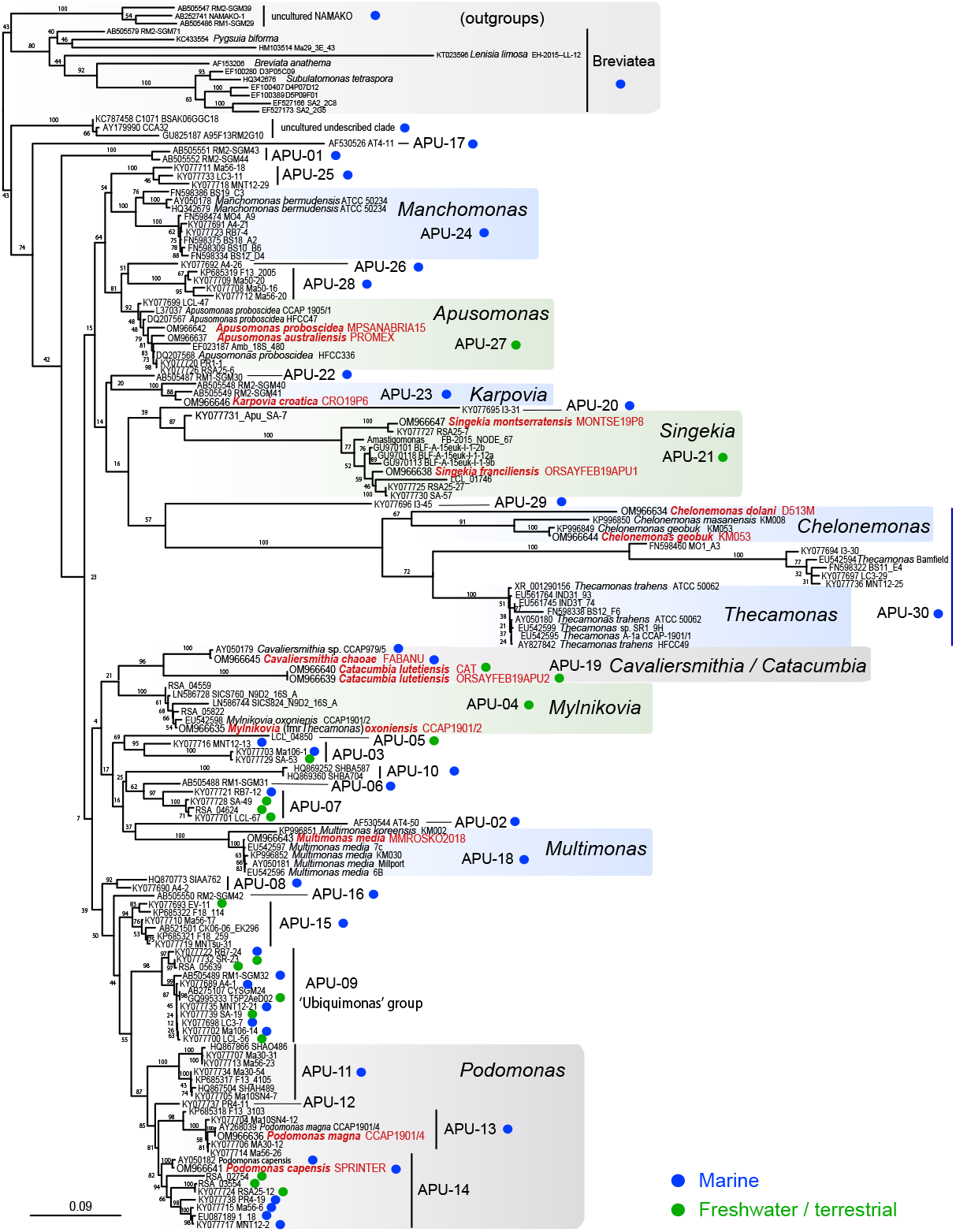
Maximum likelihood phylogenetic tree of 18S rRNA genes including all known sequences attributable to apusomonads. A total of 139 apusomonad sequences were included, together with 14 outgroup sequences from breviates and environmental samples. The tree was inferred using IQ-TREE (sequence evolution model GTR+F+R4) from a manually trimmed alignment consisting of 1,877 nucleotide positions. Support percentage values correspond to 1,000 nonparametric bootstrap replicates (bs); bullet points indicate maximum support. OTU names labeled APU-XX correspond to those identified by Torruella et al. (2017). OTU prefixes indicate environmental origin of sequences: MW, marine (blue color); FW, freshwater or soil (green color). Sequences from this study and clades representing newly described genera are indicated in bold text in red.

**Figure 2.**
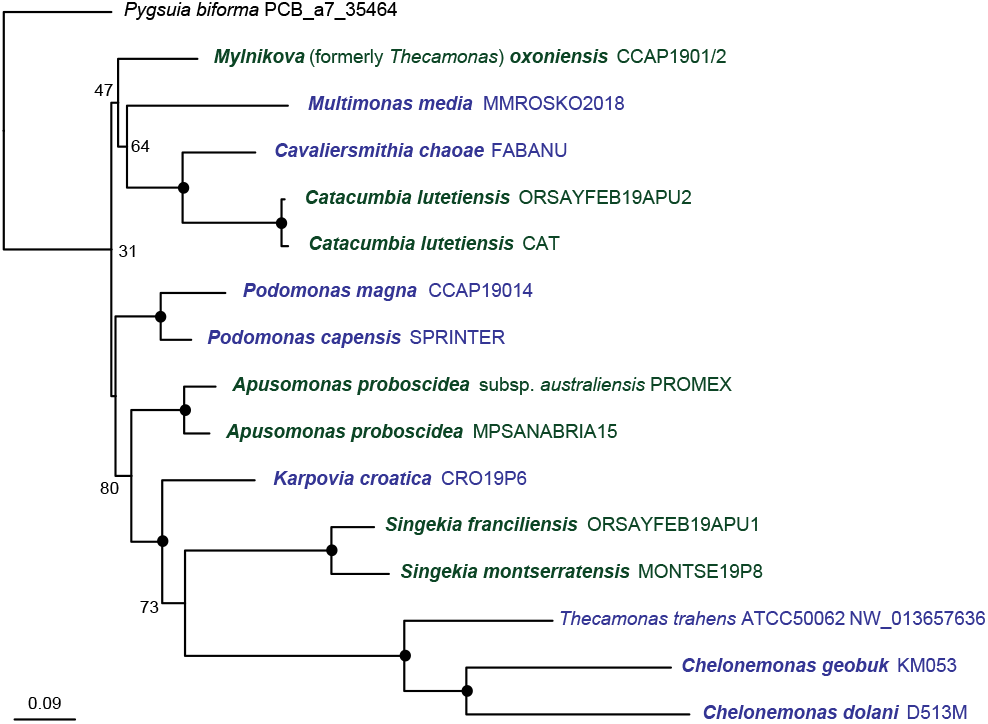
Maximum likelihood phylogenetic tree of rRNA gene operons from apusomonads examined in this study. A total of 15 apusomonad sequences were used; the breviate *Pygsuia biforma* was included as an outgroup. The tree was inferred from an automatically trimmed (trimAl) alignment of 5,596 nucleotides using IQ-TREE and the GTR+F+R4 model of sequence evolution. Support percentage values correspond to 1,000 nonparametric bootstrap replicates; black circles indicate maximum support. Sequences are colored as a function of the ecosystem type of the corresponding apusomonad species; marine (blue), freshwater/terrestrial (green). Sequences generated in this study are in bold.

### Microscopy and imaging

Optical microscopy observations were done with a Zeiss Axioplan 2 microscope equipped with oil-immersion differential interference contrast (DIC) (NEOFLUAR 100X/1.3) and phase contrast objectives. Images were taken with an AxiocamMR camera using the Zeiss AxioVision 4.8.2 SP1 suite. Videos were recorded with a Sony α9 digital camera (Supplemental Movies S1-11). Typical apusomonad morphometric data were captured through the 100x objective on 30 cells from each strain. Photos were taken at different planes to visualize different cell parts. Measurements provided are for the visible parts of the flagella, i.e., the posterior flagellar length is not measured from its point of insertion into the cell at the body’s anterior end. For measurement purposes, the proboscis sleeve length of *Apusomonas* was considered to correspond to that of the sleeve plus the portion of the mastigophore extending from the cell body flange. For all freshwater apusomonads, the measure retained for comparison was that of the widest vacuolar diameter in each cell (Table S1). For transmission electron microscopy (TEM), cell suspensions of *Multimonas media* MMROSKO2018 were mixed with an equal volume of fixative containing 2.5% glutaraldehyde and 2% paraformaldehyde in 0.2 M sodium cacodylate buffer (pH 7.2) at room temperature for 1 h. Cells were pelleted by centrifugation at 1,000 g for 5 min and rinsed with 0.2 M cacodylate buffer. Specimens were then post-fixed in 1% osmium tetroxide in cacodylate buffer at room temperature for 1 h, dehydrated through increasing ethanol series and embedded in low viscosity resin (Agar Scientific, Essex UK). Ultra-thin sections were obtained using a Leica EM UC6 ultramicrotome and double stained with uranyl acetate and lead citrate prior to their observation using a JEOL 1400 electron microscope at the Imagerie-Gif facility (Gif-sur-Yvette, France). For scanning electron microscopy (SEM), cells of *Multimonas media* MMROSKO2018 were mixed with an equal volume of fixative containing 2.5% glutaraldehyde in 0.2 M cacodylate buffer. Specimens were incubated on glass plates coated with poly-L-lysine for 2 h on ice, rinsed with 0.2 M cacodylate buffer and fixed with 1% osmium tetroxide in cacodylate buffer at room temperature for 1 h, followed by dehydration through an ethanol series. Samples were critical-point dried with CO2 using a Leica CPD300 critical point dryer, coated with platinum using a Leica ACE600 high-resolution sputter coater, and observed with a Zeiss GeminiSEM 500 field emission SEM at the Imaging Facility of the Institut de Biologie Paris Seine (Paris, France).

## RESULTS AND DISCUSSION

### Cultivation and 18S rRNA gene-based identification of new apusomonad isolates

We obtained, under aerobic conditions, eleven different mono-eukaryotic cultures of cells exhibiting morphological features typical of apusomonads. Five of the cultured strains originated from marine systems and six from continental systems (freshwater sediment or soil) worldwide (Table 1). This corroborates the cosmopolitan nature of the group and their broad ecological spectrum (Torruella et al., 2017). For comparative purposes, we also investigated four previously described apusomonad strains: *Podomonas magna* CCAP 1901/4, *Mylnikovia* (formerly *Thecamonas*) *oxoniensis* CCAP 1901/2 (Cavalier-Smith and Chao, 2010), *Chelonemonas geobuk* KM053 (Heiss et al., 2015) and *Thecamonas trahens* ATCC 50062 (Larsen and Patterson, 1990). To validate the placement of the new strains within the apusomonad clade, we first amplified and sequenced their respective 18S rRNA genes. This allowed to build a preliminary tree from which select apusomonad strains for further analysis. Subsequently, we retrieved the full rRNA operon from RNAseq data of our new selected strains to carry out more in-depth phylogenetic analysis. This analysis constituted the basis for a molecular-phylogeny-informed description and taxonomic classification of the new strains, species and genera (see below).

In addition, to place our new strains in an extended phylogenetic analysis containing a wider diversity of apusomonads, including uncultured species, we included the 18S rRNA gene sequences from the 14 newly generated operons in a larger 18S rRNA gene alignment containing both sequences of available cultured apusomonad species and environmental sequences obtained from widely diverse ecosystems (Torruella et al., 2017). The phylogenetic tree was thus inferred from a multiple alignment of 155 apusomonad sequences, including the full 18S rRNA gene sequences from our new strains, and a total of 1,877 nucleotide positions (Figure 1). In order to retain the highest possible number of aligned positions and, hence, the strongest phylogenetic signal, we limited the outgroup species to only breviates, which, with opisthokonts, are apusomonads’ closest-related lineage (Brown et al., 2013; Cavalier-Smith et al., 2014; Torruella et al., 2015), and three uncultured NAMAKO sequences related to both groups (Takishita et al., 2007).

In this specific phylogenetic tree, one group of environmental sequences retrieved from marine ecosystems segregated deeply, branching as sister to all apusomonads (including related environmental sequences; Torruella et al., 2017) with moderate (74%) support. This group was formed by two marine environmental sequences from the oxygen-depleted Cariaco basin (clone A95F13RM2G10) and the Black Sea (clone C1071 BSAK06GGC18) and a third sequence from anoxic sediments in the Great Sippewisset salt marsh (clone CCA32) (Stoeck and Epstein, 2003). Two other lineages represented only by environmental sequences branched, although without statistical support, between this lineage diverging early in the inferred tree and an unresolved larger sister group. One of these lineages (APU-17, AT4-11) corresponded to a sequence obtained from hydrothermally influenced sediment in the Mid-Atlantic Ridge (López-García et al., 2003). The other sequences (APU-1, RM2-SGM43 and 44) included two from deep-sea cold seeps (Takishita et al., 2010). Should the phylogenetic position of these uncultured groups as sister to (all other) apusomonads be confirmed, they might potentially represent lineages of anaerobic or microaerophilic protists with intermediate or assorted features between those of apusomonads and what may be their closest described group, the anaerobic or microaerophilic breviates (Brown et al., 2013; Hamann et al., 2016; Minge et al., 2009). Nonetheless, the alternative possibility that anaerobiosis/microaerophily convergently arose in putatively deep-branching apusomonads and breviates cannot be discarded at this point; culturing and characterizing both the morphology and the metabolism of those lineages would be needed in support of either hypothesis. The clade containing bona fide apusomonads included all known and newly defined genera, as well as operational taxonomic units (OTUs) previously defined from environmental sequences (Torruella et al., 2017), albeit without statistical support (23% bootstrap; Figure 1). The very recently described *Podomonas kaiyoae* branched unambiguously within APU-11 (Yabuki et al., 2022), suggesting that the genus *Podomonas* covers the OTUs APU-11 to APU-14 (Figure 1).

Since the phylogenetic signal of the 18S rRNA alignment is insufficient to resolve the internal relationships of apusomonads, we inferred a phylogenetic tree from complete ribosomal rRNA operons (i.e., the 18S, ITS1, 5.8S, ITS2, and 28S sequences) from these new strains, as well as from the only available apusomonad genome (*Thecamonas trahens* ATCC 50062 (Ruiz-Trillo et al., 2007), and the breviate *Pygsuia biforma* (Brown et al., 2013) as outgroup (Figure 2). This tree recovered a slightly different topology as compared to the 18S rRNA gene tree (Figure 1), and had higher overall statistical support, albeit deep branches were still unsupported. While in the 18S rRNA tree, *Podomonas* was sister to the clade containing *Cavaliersmithia* n. gen., *Catacumbia* n. gen., *Mylnikovia* n. comb. and *Multimonas* without support (7% bootstrap, Figure 1), in the ribosomal operon tree, *Podomonas* was sister to *Apusomonas, Karpovia* n. gen., *Singekia* n. gen., *Thecamonas* and *Chelonemonas* with 31% bootstrap (Figure 2). The most significant result shown by both trees was a clade containing the genus *Apusomonas* branching as sister to a maximally supported clade including the genera *Karpovia*, *Singekia* and the long-branching *Thecamonas* and *Chelonemonas* clade, with 15% (Figure 1) and 80% bootstrap support (Figure 2). This is the first time in which two of the five previously cultured main groups of apusomonads (the subfamilies Apusomonadinae and Thecamonadinae) have been combined in a clade with any statistical support (Heiss et al. 2017). Further, the new genera *Karpovia* and *Singekia* appeared as sister to the Thecamonadinae in both trees, with 14% bootstrap in the 18S rRNA tree (Figure 1) and with maximal support in the ribosomal operon tree (Figure 2). Meanwhile, the newly described genera *Catacumbia* and *Cavaliersmithia* formed a maximally supported clade (Figure 2), which was also well supported (96% bootstrap) in the 18S rRNA tree (Figure 1). The remaining genera, including *Podomonas*, *Multimonas*, and *Mylnikovia*, and the interrelationships of these and the abovementioned clades, remain unresolved in both trees. Fully solving apusomonad internal phylogeny will likely need phylogenomic level analyses.

### Morphological observations

We examined the morphology of our new eleven isolates together with four previously described apusomonads by light microscopy (Table 1). Observations of new isolates during active growth and late stationary phase revealed the typical fusiform *Amastigomonas*-type cell shape (see below) except for PROMEX and MPSANABRIA15, which had clearly distinguishable round *Apusomonas-type* cell shapes (Figures 3-7). Measurements of the cells are summarized in Table S1. Videos showing the movement, locomotion, cell fusion, division, and predation of these apusomonad strains can be found as Supporting Information (Movies S1-S11).

**Figure 3.**
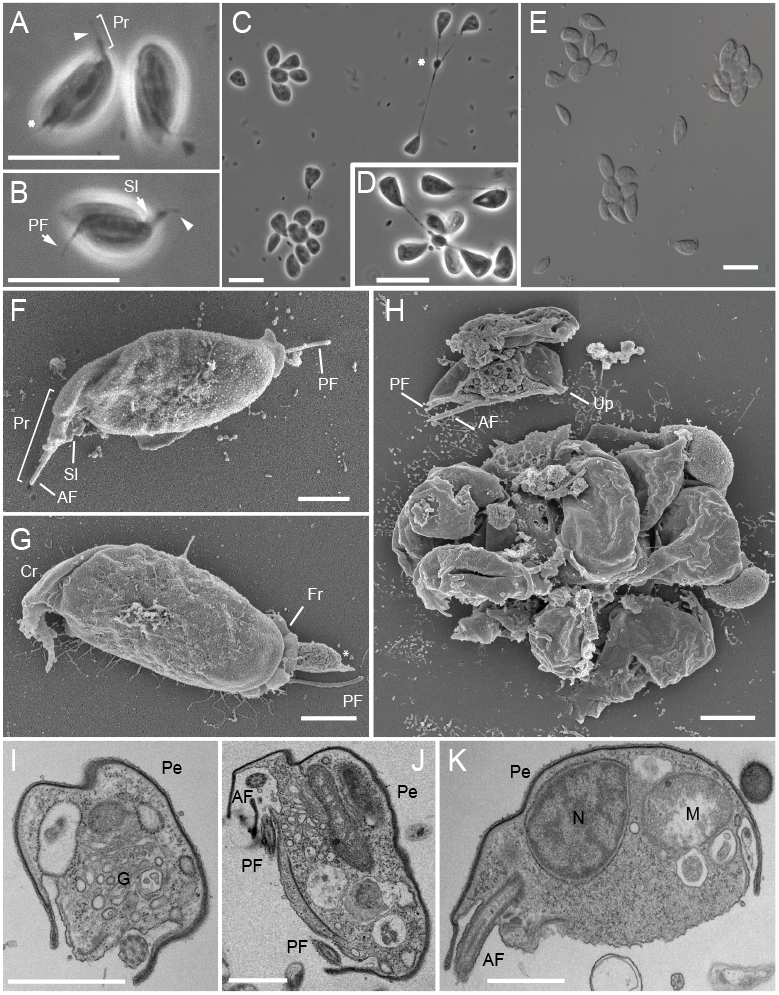
Light and electron micrographs of *Multimonas media* strain MMROSKO2018. **A-B.** Individual cells of *Multimonas media*. **C-E.** *M. media* cells dividing by multiple fission (see Movie S1). Note that the anterior part of the body does not exhibit a developed proboscis. **F-H.** Scanning electron microscopy (SEM) images showing dorsal views of the cell. **H.** SEM image of a cluster of cells possibly in multiple-fission ‘rosette’ stage. **I-K.** Transmission electron microscopy (TEM) images showing transverse (I), oblique (J), and longitudinal (K) planes of section. Note that the dorsal surface of the cell is clearly underlain by the pellicle, but not the ventral side. Scale bars: 10 μm for **A-E**; 1 μm for **F** and **G, I-K**; 2 μm for **I**. Arrowhead = acroneme; asterisk = pseudopodium; AF = anterior flagellum; Cr = anterior ‘crease’; Fr = ‘frilled’ edge of skirt; G = Golgi apparatus; M = mitochondrion; N = nucleus; Pe = pellicle; PF = posterior flagellum; Pr = proboscis; Sl = proboscis sleeve; Up = undeveloped proboscis.

All described apusomonads are gliding biflagellates (Adl et al., 2019). Their anterior flagellum runs through an open sleeve (Karpov and Zhukov, 1986), collectively termed the proboscis, which is a distinguishing feature of the group. While the ultrastructural morphology of the proboscis is distinctive, under low magnification it can be confused with the probosces of *Rhynchomonas* or certain cercozoans (Larsen and Patterson, 1990). Under TEM, all studied apusomonads possess a smooth pellicle immediately below the dorsal and lateral plasma membrane. This pellicle exhibits different ultrastructure in different lineages (Heiss et al., 2013), which may possibly be visible externally (see SEM figures in Cavalier-Smith and Chao, 2010, and Heiss et al., 2015).

Two main apusomonad cell shape types are known. One is the exclusively freshwater or soil-dwelling *‘Apusomonas-type’*, which features an ovate main cell body, with the exception of an anterior protrusion termed the mastigophore, which contains both basal bodies at its distal end, and acts as part of the proboscis (Karpov and Zhukov, 1986). The *‘Apusomonas-type’* bears a hidden posterior flagellum that trails underneath the cell and no obvious pseudopods. These cells glide in circular motion and rarely turn in angles or reverse. The other, more commonly encountered cell shape is the *Amastigomonas*-type, which has a fusiform cell body in which the lateral part of the plasma membrane projects ventrally in thin folds, forming a ‘skirt’; this skirt is underlain by the pellicle, which terminates at the skirt margin (Heiss et al., 2013, 2015). The ventral and inner lateral surfaces of the plasma membrane are relatively plastic, and can produce projections, such as lateral pseudopodia, trailing filopodia and bulbous ends, which can project visibly from the ventral and lateral surfaces of the cell (Cavalier-Smith and Chao, 2010; Myl’nikov and Myl’nikova, 2012; Heiss et al., 2013). Additionally, some species have a rigid protrusion near the base of the proboscis, a structure referred to as tusk (Heiss et al., 2013). *Amastigomonas-type* cells can contract in a circular to triangular shape to turn direction in all angles, including backwards, often by retaining a blob of cytoplasm attached to the substrate while folding the anterior part of the cell body, and after turning, they elongate again, along the direction of travel. Only freshwater species have contractile vacuoles, and to date, only one species (*Mylnikovia oxoniensis*) is known to form cysts.

All studied apusomonads can divide by longitudinal binary fission during culture growth, but some also present multiple fission of cells of equal or different size. The most characteristic step observed in *Amastigomonas*-type cells is a long cytokinesis when the daughter cells glide in opposite direction to complete the cell fission, often at a distance several times their own cell body, giving the impression that they need this mechanical stress to finish the process. Previous molecular phylogenies have suggested that the *Apusomonas*-type cell shape is derived, and that the *Amastigomonas-type* is ancestral (Cavalier-Smith and Chao, 2007; Heiss et al., 2015), a hypothesis strongly supported by our phylogeny (see above).

From the combination of molecular phylogeny and morphological characterization of our cultured apusomonad strains (Table 1 and S1), we have identified four strains of known species and seven strains representing four new genera and six new species, as follows. Although most of the newly characterized strains are of the *Amastigomonas*-type, none of them conforms to the original description of the genus *Amastigomonas* (de Saedeleer, 1931; Hamar, 1979; Zhukov, 1975; Mylnikov and Mylnikova, 2012), which is therefore not attributed to any of them.

#### Multimonas

*Multimonas media* is the type species of this marine genus (Cavalier-Smith and Chao, 2010; Heiss et al., 2015). It is a relatively large fusiform apusomonad species (~8.5 μm cell length for Cavalier-Smith’s strain; ~7.6 μm for Heiss et al.’s KM030 strain). It can divide by multiple fission, starting with a coenocyte or cytoplasmic aggregate ‘rosette’ stage containing multiple nuclei (Karpov and Mylnikov, 1989), although binary fission is also known (see below). No true cyst stage or contractile vacuole have ever been reported, and scanning electron microscopy of two strains (Heiss et al., 2015) showed no evidence of a tusk. It has been proposed that this genus was morphologically identified by Karpov and Mylnikov (1989) as *Amastigomonas caudata* or by Mylnikov (1999) as *Amastigomonas marina* (Cavalier-Smith and Chao, 2010). Only one other species of *Multimonas, M. koreensis* KM002 (~7.5 μm long), has been described; SEM imaging of both species reported probable extrusomes and a frilled or flanged margin to the ventral edge of the skirt (Heiss et al., 2015).

##### *Multimonas media* strain MMROSKO2018

This new marine strain is morphologically similar to previously studied ones. Cells are often seen advancing radially, or in an arc, from an extrapolated common point of origin, which may be the point at which multiple fission occurred (see below), grazing on bacteria and leaving the substrate clean of them behind. Cells are 6.6 ± 0.8 μm long and 3.5 ± 0.3 μm wide (ratio length/width of 1.9 ± 0.1; Figure 3A). The proboscis is obvious, with a sleeve that is 1.7 ± 0.2 μm long and an acronematic anterior flagellum projecting beyond the sleeve by 1.2 ± 0.2 μm. The posterior flagellum trails backwards through the left ventral fold and can extend behind the cell body by 2.5 ± 0.6 μm. This strain also exhibits the previously reported multiple fission (Figure 3C-E), a process we only observed from the ‘rosette’ stage (Figure 3E, H) until the end of cytokinesis. Near the end of cytokinesis in cells dividing by binary fission, cells glide in opposing directions, still attached by a thin cytoplasmic bridge that can stretch to three cell lengths long, until they mechanically detach, completing the process (Movie S1, 0:50-1:00). During this process, the cells do not appear to have a fully formed proboscis. Some cells were observed to have a round extension, separated by a constriction, from the posterior end of the cell body, especially after division (Movie S1, 0:50-1:00). Trailing filopodia can be occasionally seen under light microscopy (Figure 3A), and are obvious under SEM (Figure 3F-H). SEM images show further surface details, such the anterior ‘crease’ and the ‘frilled’ edge of the skirt. We did not observe extrusomes in our SEM preparations, unlike in (Heiss et al., 2015). We observed one possible ‘rosette’ dividing stage under SEM (Figure 3H), where cells with different sizes can be seen; although the possibility that this is an artefact cannot be excluded, the appearance of this cluster of cells is consistent with our optical microscopy observations (Figure 3C-D, movie S1). This cluster apparently includes much of the right side of a nearby, partially fragmented cell. The exposed surface of the fragment attached to the cluster has the negative impression of the material visible within the cell, and the fragment’s margins are consistent with those of the missing part of the adjacent cell. The surface of the fragment attached to the cluster extends beyond the negative impression of the cell’s inner structure, suggesting that the fragment may be continuous with the cluster; additional nearby material, both on the cell and the cluster, may or may not have been continuous as well. Regardless, it is evident that this fragmented cell is in the early stages of cytokinesis, as it has both anterior and posterior flagella trailing posteriorly (Figure 3H), the anterior flagellum being in the process of transforming into a daughter cell’s posterior flagellum. The image of this cell shows that the proboscis is not formed, since the origin of the anterior flagellum is completely naked. TEM images (Figure 3I-K) show a similar ultrastructure to other large apusomonads, with the cytoplasm containing mitochondria, dictyosomes and a circular nucleus in the anterior-dorsal part of the cell. Notice the dorsal pellicle that extends to the proboscis, and the ventral pockets, where flagella go through. Unfortunately, we did not find cells in the multiple division stage in any of our TEM sections. For extensive details on apusomonad ultrastructure, please refer to previous detailed descriptions (Karpov 1986, Molina and Nerad 1991, Karpov 2007, Mylnikov and Mylnikova 2012, Heiss et al. 2013).

#### Podomonas

The original description of this marine genus, with two species, was made by Cavalier-Smith and Chao (2010). They declared *Podomonas magna* CCAP1901/4 as the type species. Ranging from 15 to 19 μm long, this the largest apusomonad known (Cavalier-Smith et al. 2010). It has a line of refractile vacuoles on the longitudinal axis, a short sleeve in the proboscis (compared to most other fusiform apusomonads) and abundant pseudopods. Our observations of this strain suggest a tendency to float in the water column (Figure 4A), and not to glide on surfaces as do other apusomonads.

**Figure 4.**
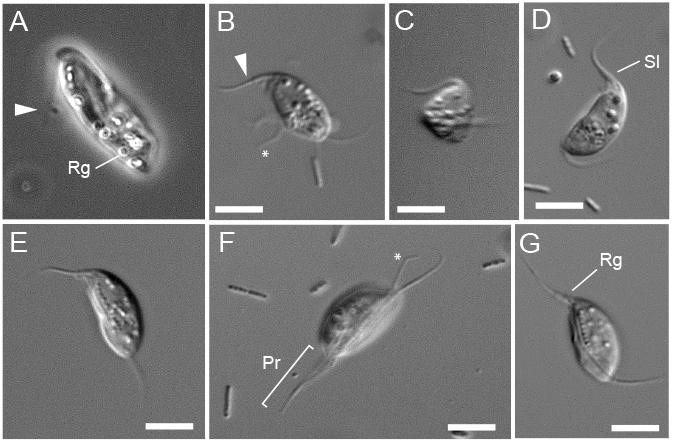
Light micrographs of living *Podomonas* species. **A.** *Podomonas magna* CCAP 1901/4, with a general view of a floating cell. **B-G.** *Podomonas capensis* strain SPRINTER. Note the abundant lateral pseudopodia and bulbous posterior end (B), that the cell folds when turning direction (C; Movie S2), the line of refractile granules (E, G), the long proboscis (F), and the conical sleeve (D). Scale bars: 5 μm. Arrowhead = acroneme; asterisk = pseudopodium; AF = anterior flagellum; PF = posterior flagellum; Pr = proboscis; Rg = refractile granules; Sl = proboscis sleeve.

*Podomonas capensis*, first described by Cavalier-Smith and Chao (2010), is more rigid than *P. magna* or any other *Amastigomonas-type* apusomonad. Fixed cells of *P. capensis* were examined under SEM and TEM, in addition to observations of live cells under light microscopy. Individual cells have a ~12 μm long cell body and comparatively long flagella (anterior, 10 μm; posterior, 14 μm). Under light microscopy, *P. capensis* was described as gliding faster than other apusomonads. It had longitudinal prominent refractile granules and irregular reticulate pseudopods. The same fixed strain was later analyzed under TEM (Figure S3 in Heiss et al. 2013), revealing the presence of a tusk, a feature shared with other apusomonads such as *Chelonemonas,Thecamonas* and the recently described marine *Podomonas kaiyoae* (Yabuki et al., 2022). *P. kaiyoae* is able to grow in axenic conditions and it is morphologically very similar to other species of the genus; interestingly, it seems to present extrusomes, like *M. media*.

##### *Podomonas capensis* strain SPRINTER

can be described in very similar terms; although cell measurements are smaller than in the no longer available strain described by Cavalier-Smith and Chao (2010), this strain was confirmed as fitting all other aspects of the species by its original describer (Cavalier-Smith, pers. comm.). The cell is 8.6 ± 0.9 μm long and 4.1 ± 0.5 μm wide (ratio of 2.1 ± 0.3). The posterior flagellum is 5.1 ± 1.2 μm long and the anterior 5.2 ± 0.7 μm long, with a sleeve of 1.6 ± 0.3 μm; this shorter sleeve compared to the whole proboscis is in agreement with the description of the genus *Podomonas* (Figure 4B-G). The dorsal pellicle appears to fold when the cell changes direction (Figure 4C; this was not described by Cavalier-Smith and Chao, 2010). Some cells exhibit bulbous posterior ends or lateral filopodia (Figure 4B); most of them present a long filopodium trailing behind the cell, parallel to the posterior flagellum (Figure 4F). Cells of this strain are more pointed at the anterior side of the cell body, due to the short, conical sleeve; glide faster than any other apusomonad in our survey; and have a prominent longitudinal line of refractile granules along the posterior flagellum (Figure 4B, E, G).

When the culture is actively growing, it is easy to see many cells floating and moving spasmodically, not attached to the surface. When attached, they glide, with slow or fast movement, while beating the anterior flagellum faster than other apusomonads (Movie S2, 0:00-0:36, 0:53-1:53). On rare occasions, we observed aggregates of cells (Movie S2, 1:28-1:34). These might be homologous to the ‘rosette’ stage of cell division by multiple fission observed in *Multimonas* and other *Amastigomonas-type* apusomonads. Usually the observed divisions were bipartitions of two cells of equal size but, occasionally, we could see smaller flagellated cells budding from a bigger cell. The small cell would appear to struggle with torsions and spasmodic movements to detach from the mother cell. After an hour, we did observe some small cells freely gliding, likely resulting from this budding process. We could not distinguish cysts. A tusk, known to exist from previous electron-microscopic characterizations (Heiss et al., 2013), was not clearly observed under light microscopy.

#### *Mylnikovia oxoniensis* n. comb

##### *Mylnikovia oxoniensis* CCAP1901/2

was originally named *Thecamonas oxoniensis*. However, our phylogenetic trees clearly placed it far from marine *Thecamonas* (Figures 1-2). Accordingly, we define a new genus, *Mylnikovia*, to accommodate this species (see nomenclatural actions below). Observations under phase contrast of the strain CCAP1901/2 confirmed previously described characteristics (Figure 5A-F), including leaf-like shaped, 9-11 μm long and 3.5-6 μm wide cells (Cavalier-Smith and Chao, 2010). The cell is elongated, with regular width up to its rounded extreme ends. It has a short anterior flagellum within a wide and short sleeve (Figure 5A). It can have a longitudinal line of refractile vacuoles (Figure 5B), as well as the typical posterior contractile vacuole of freshwater apusomonads (Figure 5C, D). Cells can display a bulbous posterior end and trailing cytoplasm (Figure 5E), but tusks were not observed. During our observations, we witnessed what appeared to be the fusion of two cells, but unfortunately we were unable to see the outcome of this process (Figure 5C, Movie S3). Multiple fission was not observed.

**Figure 5.**
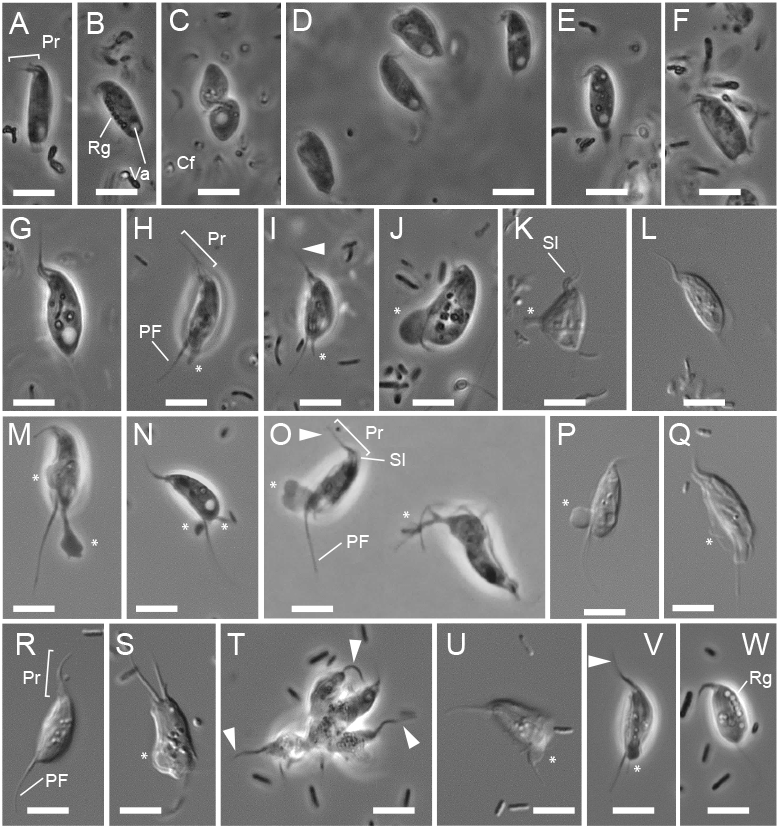
Light micrographs of *Mylnikovia* n. gen., *Catacumbia* n. gen., and *Cavaliersmithia* n. gen., **A-F.** *Mylnikovia* (formerly *Thecamonas*) *oxoniensis* CCAP 1901/2. Note the rectangular cell shape, the short flagellum inside a short and wide sleeve (A), the longitudinal line of refractile vacuoles and posterior contractile vacuole (B), the fusion of cells (C; see also Movie S3), the trailing bulbous end (E). **G-L**. *Catacumbia lutetiensis* n. sp., strain ORSAYFEB19APU2. Note the anterior flagellum inside the sleeve (G), and the trailing and lateral pseudopodia used for phagocytosis (H-K; Movie S4). **M-Q.** *Catacumbia lutetiensis* strain CAT presents abundant lateral and trailing pseudopodia (both strains can be seen in Movie S4). **R-W.** *Cavaliersmithia chaoae* n. sp., strain FABANU CCAP 1346/1 (Movie S5). Note the bicephalic ‘conjoined-twin’ stage (S), the multiple fission ‘rosette’ stage (T), and the refractile granules (W). Scale bars: 5 μm. Arrowhead = acroneme; asterisk = pseudopodium; PF = posterior flagellum; Pr = proboscis; Rg = refractile granules; Sl = proboscis sleeve.

#### *Catacumbia lutetiensis* n. gen., n. sp

##### *Catacumbia lutetiensis* strain CAT and strain ORSAYFEB19APU2

These strains of freshwater fusiform apusomonads have 99.52% identical ribosomal RNA operons, with differences occurring only in the ITS1 and ITS2 regions (Figure 2). Both strains also have short and wide sleeves and long acronematic flagella, and produce many pseudopods, like *Podomonas*. Accordingly, we classify these two strains within the same single species, although they display slightly different sizes and morphological features. Strain ORSAYFEB19APU2 (Figure 5G-L) cells are slightly bigger and rounder, being 8.0 ± 1.6 μm long and 4.4 ± 1.0 μm wide (ratio of 1.8 ± 0.3). The posterior flagellum is shorter, at 5.67 ± 2.32 μm, and the anterior is longer, at 3.9 ± 1.3 μm, with a sleeve of 1.5 ± 0.4 μm. The contractile vacuole can reach 0.8 - 2.2 μm diameter. Strain CAT (Figure 5M-Q) has cells 7.77 ± 0.74 μm long and 3.0 ± 0.5 μm wide (ratio of 2.6 ± 0.4). The posterior flagellum extends backwards by 9.4 ± 1.8 μm behind the posterior end of the cell, and the anterior, 2.7 ± 0.6 μm beyond the proboscis sleeve, which is 1.9 ± 0.4 μm long. The contractile vacuole can reach 0.8 ± 0.1 μm diameter and filopodia elongate about 3.6 ± 1.5 μm long. Strain ORSAYFEB19APU2 exhibits a leaf-shape similar to *M. oxoniensis* but is thinner at the extremes; it is also more rigid and smoother than strain CAT. Comparing their length-to-width ratios, strain CAT is thinner than ORSAYFEB19APU2. CAT also appears more flexible and lumpier. We also observed highly vacuolated cells in ORSAYFEB19APU2 more than twice the size of normal cells, which might indicate an imminent cell division. We were also able to record how a cell engulfs a bacterium, using lateral pseudopodia and retaining the food vacuole in the posterior part of the cell (Movie S4, 0:48-1:46, 4:02-4:07). Although highly vacuolated, we never observed a line of vacuoles or refractile granules as in *M. oxoniensis* or *C. chaoae*. Neither of these strains presented an obvious tusk or multiple fission.

#### *Cavaliersmithia chaoae* n. gen. n. sp

##### *Cavaliersmithia chaoae* strain FABANU

We isolated this apusomonad strain (Figure 5R-W) from a culture of the marine rhodophyte alga *Neorhodella cyanea* CCAP 1346/1, which was originally isolated from the Mediterranean Sea. Coincidentally, its 18S rRNA gene sequence is 99.61% identical to that of an uncharacterized *Amastigomonas* sp. 2 (AY050179) found as a contaminant of the marine cryptophyte alga *Cryptomonas rostrella* CCAP 979/5 from the Brittany coast (Cavalier-Smith and Chao, 2003). Unfortunately, the CCAP strain was never described or characterized, and is no longer available (Cavalier-Smith and Chao, 2010). Therefore, our morphological characterization of our strain FABANU cannot be compared to the morphology of the organism that yielded sequence AY050179.

The cell is 8.1 ± 0.8 μm long and 4.1 ± 0.5 μm wide (ratio of 1.9 ± 0.3). The posterior flagellum extends 6.9 ± 1.9 μm behind the cell, and appears more mobile than in other apusomonads. The anterior flagellum is slender, 4.1 ± 0.7 μm long, hosted within a wide and short sleeve of 1.4 ± 0.2μm. This species is highly flexible, and presents long trailing filopodia but lacks bulbous ends. It also presents a line of refractile granules, similarly to *M. oxoniensis* and *Podomonas* spp. (Figure 5W). It often presents ventral lamellipodia, similarly to *Podomonas* and *Thecamonas*. Similar to *Multimonas media*, it can divide by multiple fission in a ‘rosette’ stage (Figure 5T; Movie S5, 4:18-4:49 and 4:49-5:41). We observed many ‘conjoined-twin’ cells, with two anterior and two posterior flagella that behaved jointly as a normal cell, advancing at a similar pace and similarly whipping their anterior flagella inside a common sleeve. We could not ascertain whether this was a form of cell division or cell fusion (Movie S5, 3:44-4:18). *Catacumbia* and *Cavaliersmithia* formed a stable and robust clade in molecular phylogenetic trees (Figures 1-2) and shared relatively large cell sizes, highly vacuolated cells, proportionally shorter sleeves than most other apusomonads, and a thin anterior flagellum that can be distinguished within the sleeve under phase contrast imaging. Some of these features are also exhibited by *Mylnikovia* (Figure 5). Being genetically and morphologically very close to the freshwater *Catacumbia*, we tried to acclimate the *Cavaliersmithia* strain to freshwater by subculturing it in progressively diluted marine media, but it never grew in pure freshwater medium.

### Apusomonadinae

Karpov and Mylnikov (1989) already identified similarities between *Amastigomonas* strains and *Apusomonas*. Later, the marine *Manchomonas bermudensis* Cavalier-Smith, 2010, first described as *Amastigomonas bermudensis* (Molina and Nerad, 1991) was found to be related to the soil-dwelling *Apusomonas proboscidea* based on 18S rRNA gene trees (Cavalier-Smith and Chao, 2010). *Manchomonas bermudensis* has fusiform *Amastigomonas-type* cells, 9.5 μm long and 4 μm wide. It displays the longest known proboscis sleeve in apusomonads, almost covering the entire anterior flagellum, like its *Apusomonas* relatives. *Apusomonas proboscidea* Alexeieff, 1924 was the first apusomonad to be characterized ultrastructurally (Karpov and Zhukov, 1986) and molecularly (Cavalier-Smith and Chao, 1995).

The cell body is a rigid dorsoventrally flattened oval, almost round, with a refractive periplasm. The body contains a dense granular plasma including the posterior nucleus, the contractile vacuole, and some digestive vesicles. The proboscis is unlike that of other apusomonads in that it includes an anterior extension from the main cell body: the mastigophore, a stiff prolongation of the cell into which the flagella insert (Figure 1 in Heiss et al. 2016), and which contains both basal bodies at its distal (anterior) end (Karpov, 2007). Together with the anterior flagellum, which is visible inside the sleeve, this structure waves leftwards when viewed dorsally, while the cell glides on the substrate, usually in circular trajectories. The acroneme of the anterior flagellum is rarely visible, and the whole posterior flagellum is hidden under the mastigophore and cell body. In this work, we isolated two new strains of *Apusomonas proboscidea* and *Apusomonas australiensis*.

#### *Apusomonas proboscidea* strain MPSANABRIA15

This strain is morphologically similar to the strain CCAP 1905/1 (Cavalier-Smith and Chao, 2010). Unfortunately, we could not compare our strain with strains HFCC47 and HFCC336, since images of them have not been published (Scheckenbach et al., 2006), but previous studies point to the same morphological features. Our morphological observations are similar to those of Vickerman et al. (1974) (9 - 14 μm long, 6 - 11 μm wide), Karpov and Zhukov (1986) (8 - 12 μm) and Elekund and Patterson (1997) (7 - 8 μm); although MPSANABRIA15 displayed cells on the smaller end of the reported range: 7.9 ± 0.9 μm long and 7.9 ± 0.5 μm wide. Cells of this strain are almost perfectly round, with a length-to-width ratio of 1.0 ± 0.1 (Figure 6A-D). The uncovered anterior flagellum measured 1.3 ± 0.2 μm. Most of the proboscis consisted of the sleeve plus the mastigophore, which were together 2.6 ± 0.5 μm long. The contractile vacuole can reach 1.1 ± 0.2 μm diameter. Like other apusomonads, it glides on the substrate, although it can sometimes be seen floating with a prominent wiggling proboscis. It also divides by binary fission, with the two daughter cells gliding in opposite directions until they mechanically separate their cytoplasms (Movie S6). No filopodia were observed, in agreement with previous TEM studies (Karpov and Zhukov, 1986). However, a SEM image from an unidentified *Apusomonas* suggests radial thin filopodia emerging from the edge of the cell body (penard.de/Explorer/Apusomonadida/). It would not be too surprising for *Apusomonas* to build filopodia, given that all other apusomonads do.

**Figure 6.**
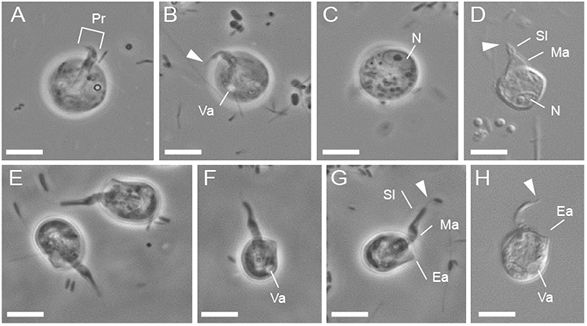
Light micrographs of living *Apusomonas*. **A-D.** *Apusomonas proboscidea* strain MPSANABRIA15 (Movie S6). Note the long mastigophore protruding from the central part of the rounded cell (A), the long sleeve and anterior flagellum (B), the contractile vacuole (B), and the large posterior nucleus with a central nucleolus (C and G). **E-H**. *Apusomonas australiensis* strain PROMEX (Movie S7). Note the anterior ‘ear’ bulge (as in Ekelund and Patterson 1997), the longer mastigophore, with a long and thick sleeve ending in a fine acronematic flagellum (E-H), and the posterior lateral contractile vacuole (F, H). Scale bars: 5 μm. Arrowhead = anterior flagellum; Ea = ‘ear’ bulge; Ma = mastigophore; N = nucleus; Pr = proboscis; Sl = proboscis sleeve; Va = contractile vacuole.

#### *Apusomonas australiensis* strain PROMEX

Alexeieff, when describing *A. proboscidea* in 1924 (10 μm long, 8 μm wide, 4 - 5 μm thick), observed asymmetry (see his Figure A, 4) similar to the *Apusomonas australiensis* described by Elekund and Patterson (1997) (6 - 10 μm long), although the early drawings make it difficult to compare. Our PROMEX strain (Figure 6E-H) clearly resembles *A. australiensis* (see Figure 2 in Elekund and Patterson 1997). It has a similar ‘ear’ bulge at the left anterior part of the body, and it is less rounded and has more conspicuous and flexible mastigophore than *A. proboscidea*. The cell body of PROMEX strain cells is 7.11 ± 0.68 μm long and 5.4 ± 0.5 μm wide, with a length/width ratio of 1.3 ± 0.1, indicating the more elongated oval form. The proportion of uncovered anterior flagellum measured 1.7 ± 0.3 μm, and the sleeve plus the protruding mastigophore, 5.3 ± 0.5 μm long. The maximal vacuole diameter was 0.7 ± 0.1 μm. No filopodia, cysts, or cells in stages of division/fusion were observed. In phylogenetic trees (Figure 1), the different strains of *Apusomonas* form a monophyletic group that includes some environmental sequences and our strains MPSANABRIA15 and PROMEX. However, the 18S rRNA gene pairwise identity between strains of this clade is relatively low (e.g. 97.1% between the two HFCC strains).

### *Singekia* n. gen

We describe two new species isolated from freshwater/continental systems for this novel genus of *Amastigomonas*-type apusomonads, *Singekia franciliensis* and *S. montserratensis*. Additionally, *Amastigomonas* sp. data from an environmental source (FB-2015, PRJNA293116) (Burki et al., 2016) can be ascribed to this genus (Figure S1). Our two new cultured strains have 92% identical 18S rRNA gene sequences, but *S. franciliensis* ORSAYFEB19APU1 has a unique insertion of 930 nucleotides long at almost the 3’ end of the sequence. Despite this molecular distance, both strains are morphologically identical, at least at optical microscopy resolution. They are ovoid cells with a small pointed bulbous end or a long trailing filopodium. The proboscis sometimes exhibits circular movement, and has a wide sleeve; the anterior flagellum is clearly visible under DIC, as in *Chelonemonas* (Figure 7). Cells also have a long posterior flagellum as compared to the cell size (Table S1), as in *Karpovia*. Unlike other apusomonads, these species do not bend much when turning, seeming more rigid. No lateral filopodia or cysts were observed.

**Figure 7.**
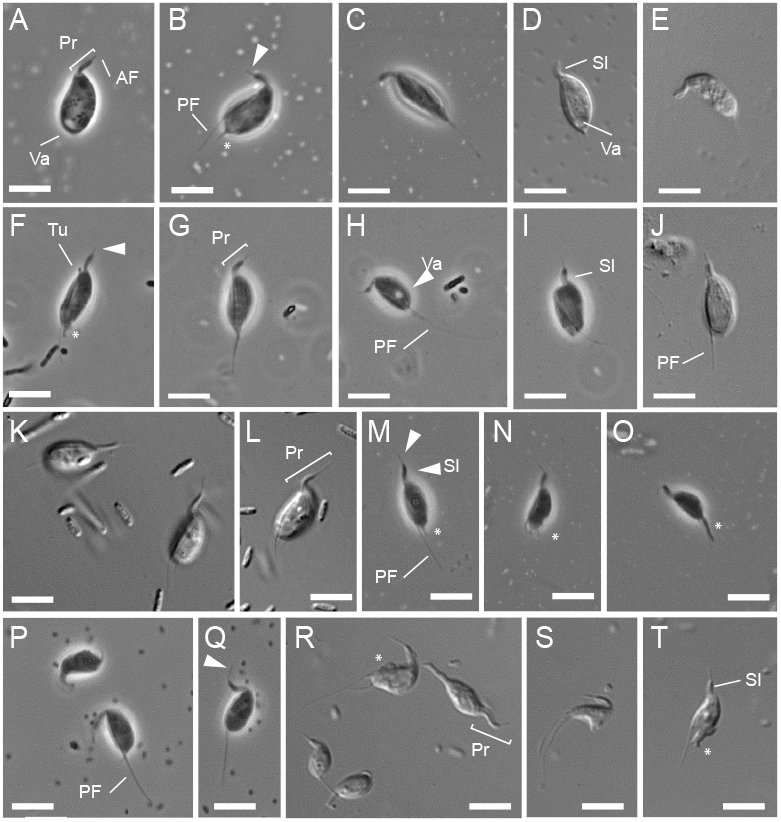
Light micrographs of species of small apusomonads belonging to the genera *Singekia* n. gen., *Karpovia* n. gen. and *Chelonemonas*. **A-E** *Singekia franciliensis* n. sp., strain ORSAYFEB19APU1 cell outline. Note the wide and long sleeve compared to the exposed anterior flagellum and the posterior vacuole (A and D). **F-J**. *Singekia montserratensis* n. sp., strain MONTSE19P8. Note the long posterior flagellum compared to the cell size, and the putative tusk (A). **K-O** *Karpovia croatica* n. sp., strain CRO19P6. Note the long proboscis and the different kinds of trailing and posterior pseudopodia (M-O). **P-T** *Chelonemonas dolani* n. sp., strain D513M cells with long sleeve and acronematic flagellum, and multiple lateral and trailing pseudopodia (R-T). Scale bars: 5 μm. Arrowhead = anterior flagellum; asterisk = pseudopodium; PF = posterior flagellum; Pr = proboscis; Sl = proboscis sleeve; Tu = Tusk; Va = contractile vacuole.

#### *Singekiafranciliensis* n. gen., n. sp., strain ORSAYFEB19APU1

Cells (Figure 7A-E) are 6.1 ± 0.6 μm long and 3.8 ± 0.3 μm wide (ratio of 1.6 ± 1.2). The posterior flagellum extends 4.5 ± 1.9 μm behind the cell, and the anterior flagellum extends 1.3 ± 0.5 μm beyond the sleeve; the sleeve is 2.1 ± 1.9 μm long. The contractile vacuole expands up to 0.7 ± 0.7 μm. This species is bigger than *S. montserratensis* (Movie S8). No tusks were observed.

#### *Singekia montserratensis* n. gen., n. sp., strain MONTSE19P8

Cells (Figure 7F-J) are 5.6 ± 0.6 μm long and 2.7 ± 0.4 μm wide (ratio of 2.1 ± 0.3). The posterior flagellum extends 3.7 ± 1 μm behind the cell, and the anterior flagellum 1.2 ± 0.5 μm beyond the proboscis sleeve, with a sleeve length of 1.7 ± 0.2 μm. The contractile vacuole expands up to 0.5 ± 0.1 μm. Some observations suggest the occurrence of division by multiple fission as in *Multimonas* or some other *Amastigomonas-type* apusomonads (Movie S9). This species seems to possess a tusk structure as described in *Podomonas*, *Chelonemonas* and *Thecamonas trahens* (Heiss et al., 2013, 2015)(Figure 7F), although this will need confirmation by electron microscopy observations.

### *Karpovia croatica* n. gen., n. sp

#### *Karpovia croatica* strain CRO19P6

This is a novel genus and species of marine *Amastigomonas-type* apusomonad. Cells (Figure 7K-O) are 5.6 ± 0.6 μm long and 3.2 ± 0.4 μm wide (ratio of 1.7 ± 0.2). The posterior flagellum extends 3.7 ± 0.9 μm behind the cell body and the anterior flagellum 1.8 ± 0.3 μm beyond the proboscis sleeve, with the sleeve being 2.2 ± 0.4 μm long. These small cells resemble those of *Singekia* and *Chelonemonas* in size and rigidity when turning direction sharply. The proboscis moves similarly to *Singekia* and thecamonadids, in more vertical or more circular movements than the flat/lateral whipping of other apusomonads; this is more evident during slow movement. Also, the sleeve is wide, as in *Singekia*. The cell often presents a trailing filopodium, and the bulbous posterior end visible when the cell turns is not as lobose as in other species, but rather presents dactylopodium-type subspseudopodia. Based on our observations, it is unclear whether this species possesses a tusk (but see Movie S10 min. 1:37). Contractile vacuoles, cysts and multiple fission were not observed.

### Thecamonadinae

This subfamily includes two genera, *Thecamonas* and *Chelonemonas*, of marine *Amastigomonas*-type apusomonads. Cells are small and display plasticity when turning, have long sleeves compared to the naked length of the anterior flagellum, and have abundant filopodia, occasionally also producing lamellipodia (Heiss et al. 2013, 2015). *Thecamonas trahens* (Larsen and Patterson, 1990) was the first *Amastigomonas-type* apusomonad to be morphologically characterized at cytoskeleton-level detail with SEM and TEM (Heiss et al., 2013), to be molecularly characterized after the determination of its 18S rRNA gene sequence (Cavalier-Smith and Chao, 2003) and to have its genome sequenced (Ruiz-Trillo et al., 2007). *Thecamonas trahens* ATCC 50062 is small, 4.5 - 7 μm long, 3 - 5 μm wide (Larsen and Patterson, 1990), and is typically seen with lateral and posteriorly trailing filopodia. Detailed TEM and SEM observations revealed a long sleeve compared to the whole proboscis, as well as the tusk (Heiss et al. 2013). The genus *Chelonemonas* originally grouped the two described species *C. geobuk* and *C. masanensis* (Heiss et al., 2015). The cells of both strains are also small (~5 and ~5.5 μm long, respectively) and display abundant lateral and trailing filopodia as well as a small tusk. SEM showed hexagonal patterns in the dorsal pellicle of both *Chelonemonas* strains (Heiss et al., 2015).

#### *Chelonemonas dolani* n. sp., strain D513M

This new species found in a mesopelagic sample (Dolan et al., 2019) exhibits cells of 4.1 ± 0.6 μm long and 2,5 ± 0.2 μm wide (ratio of 1.7 ± 0.2) (Figure 7P-T). The posterior flagellum extends 1.9 ± 5.2 μm behind the cell, and the anterior flagellum 2.1 ± 1.2 μm beyond the end of the sleeve, with the sleeve being 1.8 ± 0.5 μm long. Like *C. geobuk*, the cell is highly flexible, sometimes looking amoeboid due to the abundant lateral pseudopodia. It has a prominent sleeve, long and wide, as most small *Amastigomonas*-type apusomonads, with abundant lateral and trailing filopodia. Due to its small size, the tusk structure is difficult to observe; it seems visible in some individuals upon slight movement of the micrometer (Movie S11 at 2:22).

### Concluding remarks

We characterized several new genera and species of apusomonads, a lineage of gliding, biflagellate protists that are widely occurring, albeit in low frequency, in marine and freshwater systems worldwide. Our study extends the diversity of morphologically characterized apusomonads, including novel cultured representatives of previously defined environmental OTUs (Torruella et al., 2017), such as the marine APU-23 *Karpovia*, the freshwater APU-21 *Singekia*, and APU-19, which groups the marine *Cavaliersmithia* and the freshwater *Catacumbia*. Although rRNA-gene based phylogenetic trees do not resolve the internal topology of the apusomonad tree well, the fact that deep-branching paraphyletic subgroups of the apusomonad tree have been exclusively found in seawater environments suggests a marine origin for the clade. The placement of freshwater lineages scattered in the tree suggests multiple transitions to freshwater ecosystems (Figure 1), reinforcing previous observations (Torruella et al., 2017). Likewise, the deepest branches in the apusomonad tree correspond to environmental sequences so far only retrieved from oxygen-deprived settings, which might suggest a facultative anaerobic or microaerophilic ancestor for the clade. Despite the studied species and genera covering a large phylogenetic diversity in 18S rRNA gene trees, they share similar morphological traits with already-described apusomonads, although with some cell-size dissimilarities (Cavalier-Smith and Chao, 2003). The rounded *Apusomonas* morphotype seems to have evolved from a fusiform *Amastigomonas*-type ancestor, likely with a large sleeve as in *Apusomonas’*s sister taxon, *Manchomonas bermudensis*, which would seem more adapted to soils (Cavalier-Smith and Chao, 2010). Therefore, the ancestor of the whole apusomonad clade likely was a fusiform biflagellate with a dorsal pellicle, a ventral cell body bounded only by the plasma membrane, and a sleeve covering only part of the anterior flagellum. We also observed stages of division by multiple fission in several apusomonad species. These observations, if confirmed and extended to other apusomonads, might suggest that division by multiple fission might have also been present in the apusomonad ancestor. They may have originated earlier, since they are also present in both lineages most closely related to apusomonads: the breviates (Walker et al., 2006) and the opisthokonts, the latter group having given rise to, among others, fungi and animals. As in many amoebozoans (Kang et al., 2017) and opisthokont protists (Dayel et al., 2011), the apusomonad life cycle appears complex, having more than a simple asexual cell division phase, as attested by the ‘rosette’ stage and multiple fission observed in *Multimonas media*, *Cavaliersmithia chaoae*, *Singekia montserratensis* and *Podomonas capensis*. Although we witnessed what might potentially be cell fusion only in *Mylnikovia oxoniensis* (Movie S3), we observed morphological and behavioral diversity of cell-types in each culture, including floating cells in most species, small cells attached to big ones in *Podomonas capensis* (Movie S2) or ‘conjoined twins’ in *Cavaliersmithia chaoae*, highly suggestive of sex. In *P. capensis*, the observation of budding cells from a larger mother cell might suggest either gamete production or multiple rounds of mitotic division from the mother cell. Additional future TEM and SEM analyses will help to characterize the cell ultrastructure of the different apusomonad species better, and may reveal morphological features, in agreement with the phylogenetic position of the different apusomonad taxa, that will provide additional morphological identities for members of this superficially conservative group. Future phylogenomic analyses based on genomics and/or transcriptomics of the different species will help to resolve the internal phylogeny of the clade, and should provide a solid basis to understand the evolution of this still poorly studied, but evolutionarily significant, lineage.

## TAXONOMIC SUMMARY

All taxonomic descriptions in this work were approved by all authors.

Eukarya: Amorphea: Obazoa: Apusomonadida Karpov and Mylnikov 1989

*Mylnikovia* Torruella, Galindo et al., n. gen.

**Diagnosis.** As for type species (originally *Thecamonas oxoniensis* Cavalier-Smith and Chao 2010).
**Etymology.** The name *Mylnikovia* is to honor the work of professor Alexander Mylnikov on the discovery and description of several *Amastigomonas*-type apusomonads, as well as his work on the first ultrastructural studies of the group. *Mylnikovia* is a first-declension feminine Latin noun.
**Type species.** *Mylnikovia oxoniensis* n. comb.

*Mylnikovia oxoniensis* Torruella, Galindo et al., n. comb.

**Diagnosis.** See Cavalier-Smith and Chao 2010.
**Type material.** See Cavalier-Smith and Chao 2010.
**Gene sequence.** The full ribosomal RNA operon sequence from *Mylnikovia oxoniensis* (strain IVY8c) was deposited in GenBank with accession number OM966635.
**Zoobank registration.** urn:lsid:zoobank.org:act:AAAEA83A-82CC-4F20-8B90-7009B605DB96

*Catacumbia* Torruella, Galindo et al., n. gen.

**Diagnosis.** As for type species, *Catacumbia lutetiensis*, Torruella, Galindo et al., this work.
**Etymology.** The name *Catacumbia* refers to the catacombs of Paris from where the sample containing this strain was obtained. *Catacumbia* is a first-declension feminine Latin noun.
**Zoobank registration.** urn:lsid:zoobank.org:act:52DE3766-EDE7-4E70-8ED2-90543403601B

*Catacumbia lutetiensis* Torruella, Galindo et al., n. sp.

**Diagnosis.** Fresh-water *Amastigomonas*-type cell with a short, wide sleeve and long acronematic anterior flagellum. Posterior flagellum long and conspicuous. Abundant lateral and trailing filopodia, abundant small vacuoles, posterior contractile vacuole. At least two cultured morphotypes with no genetic differences.
**Type culture.** A cryopreserved culture is deposited in the culture collection of DEEM laboratory, CNRS and Université Paris-Saclay (France) as the type strain CAT.
**Type locality.** Specimen isolated from freshwater sediments of the catacombs of Paris, France.
**Isolator:** Guifré Torruella isolated strain CAT and Luis Javier Galindo isolated ORSAYFEB19APU2.
**Etymology.** The name *lutetiensis* refers to the Roman city of Lutetia, which was the predecessor of the modern-day city of Paris, where the sample was collected.
**Gene sequence.** The full ribosomal RNA operon sequences from *Catacumbia lutetiensis* (strain CAT [type sequence] and ORSAYFEB19APU2) were deposited in GenBank with accession numbers OM966640 and OM966639, respectively.
**Zoobank registration.** urn:lsid:zoobank.org:act:1D29B44F-EA8F-48C2-984E-508DEBE3C409

*Cavaliersmithia* Torruella, Galindo et al., n. gen.

**Diagnosis.** As for type species, *Cavaliersmithia chaoae*, Torruella, Galindo et al., this work.
**Etymology.** The name *Cavaliersmithia* is to honor the work of Thomas Cavalier-Smith on the discovery and description of several *Amastigomonas-type* apusomonads; he was also the first to suggest the group’s sister relationship to opisthokonts, and to emphasize its evolutionary importance. *Cavaliersmithia* is a first-declension feminine Latin noun.
**Zoobank registration.** urn:lsid:zoobank.org:act:2CBA9250-1C88-411D-AA23-CBFF35C27E62

*Cavaliersmithia chaoae* Torruella, Galindo et al., n. sp.

**Diagnosis.** Marine *Amastigomonas*-type cell with short, wide sleeve and long acronematic anterior flagellum. Posterior flagellum long and conspicuous. Presents few lateral and trailing filopodia, abundant small vacuoles and refractile granules. Cells can present stages such as bicephalic ‘conjoined-twin’ and multiple fission in ‘rosette’.
**Type culture.** A cryopreserved culture is deposited in the culture collection of DEEM laboratory, CNRS and Université Paris-Saclay (France) as the type strain FABANU.
**Isolator:** Guifré Torruella.
**Type locality.** Specimen isolated from the *Neorhodella* culture CCAP 1346/1, originally from marine water from Menton, France.
**Etymology.** The name *chaoae* is to honor the work of Ema Chao on the discovery and description of several *Amastigomonas-type* apusomonads.
**Gene sequence.** The full ribosomal RNA operon sequence from *Cavaliersmithia chaoae* (strain FABANU) was deposited in GenBank with accession number OM966645.
**Zoobank registration.** urn:lsid:zoobank.org:act:C47D366F-234F-4093-8987-FCBB11EBF9AB

*Singekia* Torruella, Galindo et al., n. gen.

**Diagnosis.** Fresh-water *Amastigomonas*-type cell with a long, tight sleeve and short acronematic anterior flagellum. Posterior flagellum may or may not be conspicuous. Cells are small and rigid, presenting a posterior contractile vacuole, with small pointed bulbous end or a long trailing filopodium. Two cryptic species with identical morphology, distinguished by 18S ribosomal RNA gene sequence.
**Etymology.** The name *Singekia* refers to the SINGEK consortium, an EU H2020 Marie Skłodowska-Curie Innovative Training Network in which L.J.G performed his Ph.D. research. *Singekia* is a first-declension feminine Latin noun.
**Zoobank registration.** urn:lsid:zoobank.org:act:A3E33D03-1FDA-4B2C-B503-DD047B3FA11C

*Singekia montserratensis* Torruella, Galindo et al., n. sp.

**Diagnosis.** As for type genus, *Singekia*, Torruella, Galindo et al., this work.
**Type culture.** A cryopreserved culture is deposited in the culture collection of DEEM laboratory, CNRS and Université Paris-Saclay (France) as the type strain MONTSE19P8.
**Isolator:** Luis Javier Galindo.
**Type locality.** Specimen isolated from a puddle of rainwater in the Montserrat mountain, Catalonia, Spain.
**Etymology.** The name *montserratensis* refers to the sample obtained during a hike in Montserrat mountain with members of the SINGEK consortium.
**Gene sequence.** The full ribosomal RNA operon sequence from *Singekia montserratensis* (strain MONTSE19P8) was deposited in GenBank with accession number OM966647.
**Zoobank registration.** urn:lsid:zoobank.org:act:C8A8B20A-D240-4875-A99D-1ED487FAE9DF

*Singekia franciliensis* Torruella, Galindo et al., n. sp.

**Diagnosis.** As for type genus, *Singekia*, Torruella, Galindo et al., this work. The 18S rRNA gene sequence contains an insertion of 930 nucleotides long close to the 3’ end of the sequence.
**Type culture.** A cryopreserved culture is deposited in the culture collection of DEEM laboratory, CNRS and Université Paris-Saclay (France) as the type strain ORSAYFEB19APU1.
**Isolator:** Luis Javier Galindo.
**Type locality.** Specimen isolated from sediments from a street gutter in Orsay, France.
**Etymology.** The name *franciliensis* refers to the region of origin of the sample (Ile-de-France) from which the organism was isolated from.
**Gene sequence.** The full ribosomal RNA operon sequence from *Singekia franciliensis* (strain ORSAYFEB19APU1) was deposited in GenBank with accession number OM966638.
**Zoobank registration.** urn:lsid:zoobank.org:act:0FEAF9FA-6DE7-49B9-A51B-4D661BB6654D

*Karpovia* Torruella, Galindo et al., n. gen.

**Diagnosis.** As for type species, *Karpovia croatica*, Torruella, Galindo et al., this work.
**Etymology.** The name *Karpovia* is to honor the work of Sergey Karpov on the discovery and description of several *Amastigomonas*-type apusomonads as well as his pioneering work in apusomonad ultrastructural studies. *Karpovia* is a first-declension feminine Latin noun.
**Zoobank registration.** urn:lsid:zoobank.org:act:3F8F4CC7-B2C1-4117-9AB0-F577674B3C3B

*Karpovia croatica* Torruella, Galindo et al., n. sp.

**Diagnosis.** Marine *Amastigomonas*-type cell with long, tight sleeve and short acronematic anterior flagellum. Posterior flagellum may or may not be conspicuous. Cells small and rigid, with trailing dactylopodia or filopodia.
**Type culture.** A cryopreserved culture is deposited in the culture collection of DEEM laboratory, CNRS and Université Paris-Saclay (France) as the type strain CRO19P6.
**Isolator:** Luis Javier Galindo.
**Type locality.** Specimen isolated from the sediments of the marine lake Malo jezero in the island of Mljet, Croatia.
**Etymology.** The name *croatica* refers to the country of origin of the sample (Croatia).
**Gene sequence.** The full ribosomal RNA operon sequence from *Karpovia croatica* (strain CRO19P6) was deposited in GenBank with accession number OM966646.
**Zoobank registration. urn:lsid:zoobank.org:act:669C028C-9725-4B3C-ADCC-4EC381EA8F92**

*Chelonemonas dolani* Torruella, Galindo et al., n. sp.

**Diagnosis.** Marine *Amastigomonas*-type cell with long, tight sleeve and short acronematic anterior flagellum. Conspicuous long posterior flagellum. Cells small and flexible, presenting lateral pseudopodia.
**Type culture.** A cryopreserved culture is deposited in the culture collection of DEEM laboratory, CNRS and Université Paris-Saclay (France) as the type strain D513M.
**Isolator:** Maria Ciobanu.
**Type locality.** Specimen isolated from a coastal Mediterranean sample at Villefranche-sur-Mer, France.
**Etymology.** The name *dolani* honors work on marine protists by John R. Dolan at Villefranche-sur-Mer.
**Gene sequence.** The full ribosomal RNA operon sequence from *Chelonemonas dolani* (strain D513M) was deposited in GenBank with accession number OM966634.
**Zoobank registration.** urn:lsid:zoobank.org:act:7CD10382-A329-4D86-AD76-7E7F66443DB2

## ACKNOWLEDGMENTS

The authors acknowledge the importance of previous studies on these protists by several researchers and thank T. Cavalier-Smith, E. Chao, D. Patterson, A. Mylnikov, and S. A. Karpov for their legacy and, in specific cases, useful discussion and collaboration. We thank J. Dolan for access to samples from Villefranche-sur-Mer from which *C. dolani* was cultured, A. Guillén, author of ‘Proyecto agua’, for sending samples from the Sanabria glacial lake, and J. Favate for collecting the material from which SPRINTER was isolated. We also thank Kaleigh Lukacs for assisting with culture maintenance. This work was supported by the European Research Council (ERC) Advanced Grants ‘Protistworld’ and ‘Plast-Evol’ (322669 and 787904, respectively) and the Horizon 2020 research and innovation program under the Marie Skłodowska-Curie ITN project SINGEK H2020-MSCA-ITN-2015-675752 (http://www.singek.eu/). GT was supported by the 2019 BP 00208 Beatriu de Pinos-3 Postdoctoral Program (BP3; 801370).

## AUTHOR CONTRIBUTION

PLG, DM, GT and LG designed the study. GT, LG, MC and AH carried out isolations and laboratory work. NY performed electron microscopy studies. AH contributed strains for comparison. PLG and DM obtained funding to carry out the work. GT wrote the preliminary draft of the manuscript. PLG wrote the final manuscript. All authors read and commented on the manuscript.

## SUPPORTING INFORMATION

**Movie S1.** Video clip of *Multimonas media* strain MMROSKO2018. https://youtu.be/FHptgPcV784

**Movie S2.** Video clip of *Podomonas capensis* strain SPRINTER. https://youtu.be/Aj9V8PC66x8

**Movie S3.** Time-lapse of *Mylnikovia* (formerly *Thecamonas) oxoniensis* cells fusing. https://youtu.be/CSPTqPsYloE

**Movie S4.** Video clip of *Catacumbia lutetiensis* strain CAT and strain ORSAYFEB19APU2. https://youtu.be/HAkJvqWXXGM

**Movie S5.** Video clip of *Cavaliersmithia chaoae* FABANU. https://youtu.be/P9FXY_5XkPU

**Movie S6.** Video clip of *Apusomonas proboscidea* strain MPSANABRIA15. https://youtu.be/0vDBMmRS4l0

**Movie S7.** Video clip of *Apusomonas australiensis* strain PROMEX. https://youtu.be/pjxs7v9MCfc

**Movie S8.** Video clip of *Singekia franciliensis* strain ORSAYFEB19APU1. https://youtu.be/-VeFZ-2r3Bo

**Movie S9.** Video clip of *Singekia montserratensis* strain MONTSE19P8. https://youtu.be/_2R8rd5TPEk

**Movie S10.** Video clip of *Karpovia croatica* strain CRO19P6. https://youtu.be/QN03vdoexIo

**Movie S11.** Video clip of *Chelonemonas dolani* strain D513M. https://youtu.be/Q_03Gc6q1S0

All the videos can be accessed at once from the DEEM team YouTube channel: https://www.youtube.com/channel/UCIYNzs3VyCa-Xno121Z2cSQ

**Table S1.**
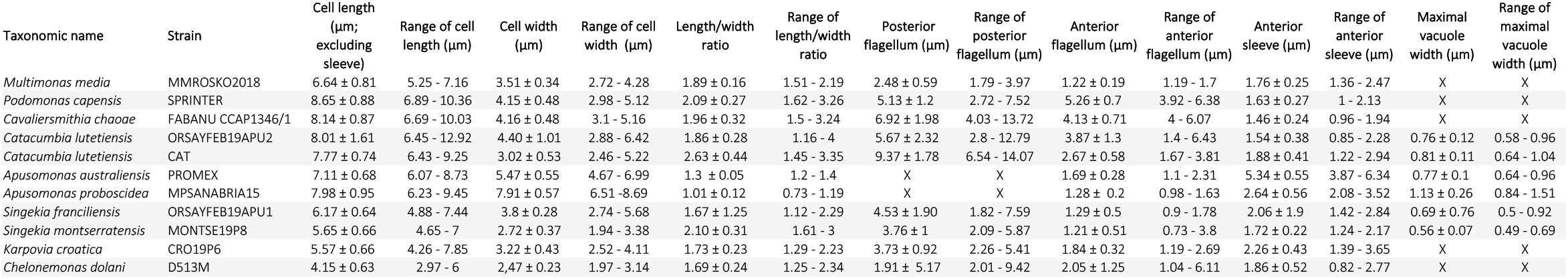
Morphometric characteristics of the studied strains by light microscopy. Cell length, width, length/width ration and length of posterior flagellum represent the average with standard deviation. For Apusomonas strains, the mastigophore and sleeve’s length are measured together. ‘Vacuole’ refers to contractile vacuoles.

| Taxonomic name | Strain | Cell length<br>(μm;<br>excluding<br>sleeve) | Range of cell<br>length (μm) | Cell width<br>(μm) | Range of cell<br>width (μm) | Length/width<br>ratio | Range of<br>length/width<br>ratio | Posterior<br>flagellum (μm) | Range of<br>posterior<br>flagellum (μm) | Anterior<br>flagellum (μm) | Range of<br>anterior<br>flagellum (μm) | Anterior<br>sleeve (μm) | Range of<br>anterior<br>sleeve (μm) | Maximal<br>vacuole<br>width (μm) | Range of<br>maximal<br>vacuole<br>width (μm) |
| --- | --- | --- | --- | --- | --- | --- | --- | --- | --- | --- | --- | --- | --- | --- | --- |
| *Multimonas media* | MMROSKO2018 | 6.64 ± 0.81 | 5.25 - 7.16 | 3.51 ± 0.34 | 2.72 - 4.28 | 1.89 ± 0.16 | 1.51 - 2.19 | 2.48 ± 0.59 | 1.79 - 3.97 | 1.22 ± 0.19 | 1.19 - 1.7 | 1.76 ± 0.25 | 1.36 - 2.47 | X | X |
| *Podomonas capensis* | SPRINTER | 8.65 ± 0.88 | 6.89 - 10.36 | 4.15 ± 0.48 | 2.98 - 5.12 | 2.09 ± 0.27 | 1.62 - 3.26 | 5.13 ± 1.2 | 2.72 - 7.52 | 5.26 ± 0.7 | 3.92 - 6.38 | 1.63 ± 0.27 | 1 - 2.13 | X | X |
| *Cavaliersmithia chaoae* | FABANU CCAP1346/1 | 8.14 ± 0.87 | 6.69 - 10.03 | 4.16 ± 0.48 | 3.1 - 5.16 | 1.96 ± 0.32 | 1.5 - 3.24 | 6.92 ± 1.98 | 4.03 - 13.72 | 4.13 ± 0.71 | 4 - 6.07 | 1.46 ± 0.24 | 0.96 - 1.94 | X | X |
| *Catacumbia lutetiensis* | ORSAYFEB19APU2 | 8.01 ± 1.61 | 6.45 - 12.92 | 4.40 ± 1.01 | 2.88 - 6.42 | 1.86 ± 0.28 | 1.16 - 4 | 5.67 ± 2.32 | 2.8 - 12.79 | 3.87 ± 1.3 | 1.4 - 6.43 | 1.54 ± 0.38 | 0.85 - 2.28 | 0.76 ± 0.12 | 0.58 - 0.96 |
| *Catacumbia lutetiensis* | CAT | 7.77 ± 0.74 | 6.43 - 9.25 | 3.02 ± 0.53 | 2.46 - 5.22 | 2.63 ± 0.44 | 1.45 - 3.35 | 9.37 ± 1.78 | 6.54 - 14.07 | 2.67 ± 0.58 | 1.67 - 3.81 | 1.88 ± 0.41 | 1.22 - 2.94 | 0.81 ± 0.11 | 0.64 - 1.04 |
| *Apusomonas australiensis* | PROMEX | 7.11 ± 0.68 | 6.07 - 8.73 | 5.47 ± 0.55 | 4.67 - 6.99 | 1.3 ± 0.05 | 1.2 - 1.4 | X | X | 1.69 ± 0.28 | 1.1 - 2.31 | 5.34 ± 0.55 | 3.87 - 6.34 | 0.77 ± 0.1 | 0.64 - 0.96 |
| *Apusomonas proboscidea* | MPSANABRIA15 | 7.98 ± 0.95 | 6.23 - 9.45 | 7.91 ± 0.57 | 6.51 - 8.69 | 1.01 ± 0.12 | 0.73 - 1.19 | X | X | 1.28 ± 0.2 | 0.98 - 1.63 | 2.64 ± 0.56 | 2.08 - 3.52 | 1.13 ± 0.26 | 0.84 - 1.51 |
| *Singekia franciliensis* | ORSAYFEB19APU1 | 6.17 ± 0.64 | 4.88 - 7.44 | 3.8 ± 0.28 | 2.74 - 5.68 | 1.67 ± 1.25 | 1.12 - 2.29 | 4.53 ± 1.90 | 1.82 - 7.59 | 1.29 ± 0.5 | 0.9 - 1.78 | 2.06 ± 1.9 | 1.42 - 2.84 | 0.69 ± 0.76 | 0.5 - 0.92 |
| *Singekia montserratensis* | MONTSE19P8 | 5.65 ± 0.66 | 4.65 - 7 | 2.72 ± 0.37 | 1.94 - 3.38 | 2.10 ± 0.31 | 1.61 - 3 | 3.76 ± 1 | 2.09 - 5.87 | 1.21 ± 0.51 | 0.73 - 3.8 | 1.72 ± 0.22 | 1.24 - 2.17 | 0.56 ± 0.07 | 0.49 - 0.69 |
| *Karpovia croatica* | CRO19P6 | 5.57 ± 0.66 | 4.26 - 7.85 | 3.22 ± 0.43 | 2.52 - 4.11 | 1.73 ± 0.23 | 1.29 - 2.23 | 3.73 ± 0.92 | 2.26 - 5.41 | 1.84 ± 0.32 | 1.19 - 2.69 | 2.26 ± 0.43 | 1.39 - 3.65 | X | X |
| *Chelonemonas dolani* | D513M | 4.15 ± 0.63 | 2.97 - 6 | 2,47 ± 0.23 | 1.97 - 3.14 | 1.69 ± 0.24 | 1.25 - 2.34 | 1.91 ± 5.17 | 2.01 - 9.42 | 2.05 ± 1.25 | 1.04 - 6.11 | 1.86 ± 0.52 | 0.82 - 2.77 | X | X |

## Notes

### Competing Interest Statement

The authors have declared no competing interest.

